# The individuality of single-frame functional brain connectivity

**DOI:** 10.64898/2026.01.05.675158

**Authors:** Clayton C. McIntyre, Heather M. Shappell, Mohsen Bahrami, Robert G. Lyday, Paul J. Laurienti

## Abstract

Converging evidence from studies on brain network “fingerprinting” and precision functional mapping suggest that brain networks are highly individualized in functionally meaningful ways. Concurrently with a growth in studies on this topic, there has been a rise in interest on dynamics (approximately second-to-second changes) in brain networks within scan sessions. While analyses of traditional static networks have increasingly grown towards emphasizing the importance of individual differences in brain network topology, studies of dynamic networks typically follow methodology that require brain states to be considered at a group level. Recent studies have begun to assess the individuality of recurring dynamic brain “states”. In this work, we extend this recent work by exploring the extent to which functional connectivity fingerprinting is feasible at single-frame temporal resolution. We estimate connectivity at individual volumes using phase coherence. We find that the identity of participants can be classified based on single volumes given sufficient database scan data and that having more highly parcellated atlases facilitates identification. Finally, we find that tasks can be identified more readily within subjects than between subjects. We conclude that participant identity may be an important driver of observed single-volume connectivity patterns. Further, the single-volume neural correlates of a task appear to be more consistent within subjects than between subjects. This highlights the importance of considering individual variability in studies of brain network dynamics.

## 1. Introduction

Our understanding of the human brain has been revolutionized by network analyses of functional neuroimaging data. Under the “network neuroscience” paradigm, the brain is modeled as a collection of nodes (representing brain regions) and edges (representing functional connections between brain regions)(Bullmore & Sporns, 2009). Functional connectivity is most commonly assessed by the synchrony of blood-oxygen-level-dependent (BOLD) signal fluctuations in functional magnetic resonance imaging (fMRI)(Ogawa et al., 1990). Traditionally, network neuroscience studies have sought to identify commonalities in brain networks across groups of people. This approach has led to the discovery of benchmarks of typical functional connectivity and the ability to relate divergence from typical connectivity patterns to neuropsychiatric or cognitive outcomes (Zhang et al., 2021). While these discoveries have advanced our understanding of the brain’s functional organization at a population level, focusing on commonalities across people often results in overlooking individual differences in brain organization that may be deeply meaningful.

Substantial effort has gone towards empirically demonstrating that individual differences in brain network topology are robust and functionally significant. For example, it has been shown that brain networks effectively serve as “fingerprints” such that an individual’s brain network can be reliably identified out of a large sample (Anderson et al., 2011; Finn et al., 2015). Further, the connectivity differences that best distinguish between people are also related to intelligence (Finn et al., 2015) and can distinguish between tasks (Anderson et al., 2011). In addition to fingerprinting, “precision functional mapping” has become increasingly popular in the literature (Gordon et al., 2017). Proponents of precision functional mapping endorse collecting large quantities of scan data in smaller samples rather than collecting limited scan data from a large sample. This practice has been used to demonstrate that while there are overarching similarities in functional networks across people, each person’s network is highly unique.

Concurrently with growing interest in the individuality of static brain networks, there has been a drastic increase in studies on dynamic brain networks (Bassett & Sporns, 2017; Lurie et al., 2020). Studies in this area examine within-session changes in brain network topology and how they relate to cognition, behavior, and neuropsychiatric disease (Allen et al., 2014; Chang & Glover, 2010; Handwerker et al., 2012). This research is motivated by the idea that traditional static networks provide useful representations of the brain’s average functional connectivity but overlook meaningful moment-to-moment variability. Despite substantial interest in dynamic brain networks, meaningful interpretation of brain network dynamics using fMRI data has proven to be a formidable challenge.

In efforts to address this challenge, many methodologies have been developed. While they differ in several ways, the most popular approaches for studying brain network dynamics have very similar analysis structures (Allen et al., 2014; Cabral et al., 2017; Shappell et al., 2019; Vidaurre et al., 2018). That is, the approaches cited above first identify a small number of functional connectivity “states” that are shared across a sample. After identifying group-level states, state sequences are inferred for each individual. Analyses are then focused on metrics of the state sequences. Because states are shared at the group level, this approach allows direct comparison of the dynamics of individuals or groups. However, this approach also forces all analyses to be carried out from a group-level perspective. In other words, a participant’s trajectory through states can be unique – but the states themselves cannot. Like early analyses of static networks, this practice emphasizes the importance of the group over the individual. It may also suffer from the same pitfall of overlooking meaningful individuality. While static network analyses have increasingly grown towards studying individuals with high precision, analyses of dynamic networks have largely been limited to group-level questions.

Recognizing this potential limitation, several studies have begun to investigate the extent to which “states” are unique to individuals. Prior work has used sliding-window connectivity (which captures average connectivity over several volumes) to evaluate the fingerprinting capability of matrices representing the variability around a static network over time (Liu et al., 2018), of a small set of “states” identified by within-subject clustering algorithms (Ghaffari et al., 2025; Menon & Krishnamurthy, 2019), and of windows of different lengths (Van De Ville et al., 2021). Additionally, edge-based time series (Faskowitz et al., 2020) have been used to estimate BOLD cofluctuations at single volumes before binning those volumes into groups to use for fingerprinting (similar to the clustering approach described above with sliding window)(Cutts et al., 2023).

To our knowledge, no study has assessed the extent to which connectivity estimates at single-volume resolution can be used for fingerprinting. Therefore, in this work we use phase coherence (Cabral et al., 2017; Glerean et al., 2012) - a measure that estimates synchrony of BOLD phases between each pair of nodes at each fMRI scan volume – to attempt to identify individuals. We investigate single-frame dynamic fingerprinting feasibility in three independent samples. We also investigate the extent to which atlas parcellation and amount of scan data influence identification accuracy. Finally, we determine whether our single-frame dynamic fingerprinting approach can identify the task a participant is doing and whether single-frame connectivity during tasks differ between participants.

## 2. Methods

### 2.1 Samples and Data Collection

Data from three existing studies were used for analyses in this work. An overview of the types of scans collected and the timing of scanning visits for each of the studies are depicted in Figure 1.

**Figure 1.**
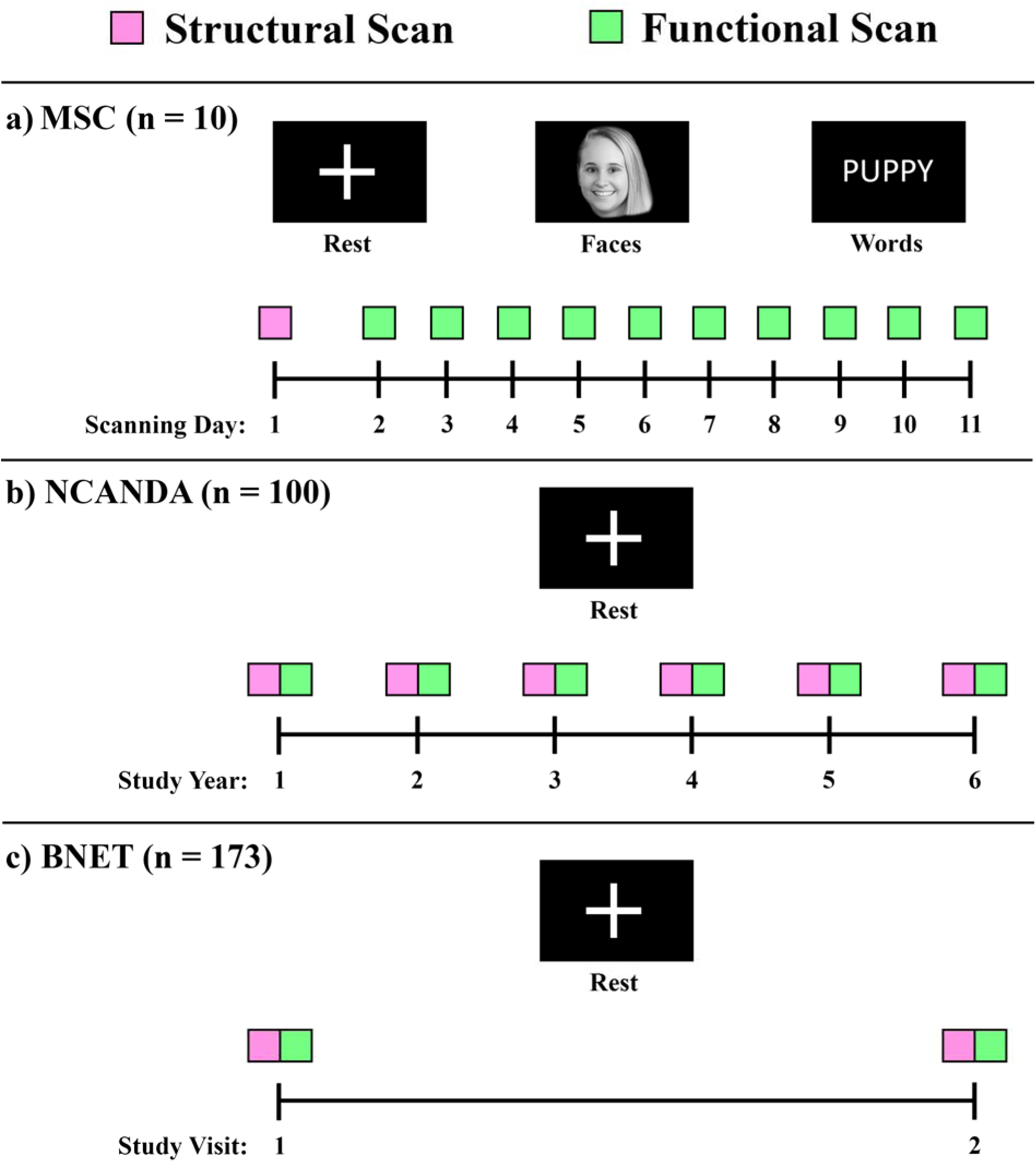
Panel **a)** depicts scan collection for the Midnight Scan Club (MSC). Structural scanning was completed before functional scanning on a separate day. Resting-state and task-based functional scans were completed over ten approximately consecutive days. To avoid copyright infringement, the face in the “Faces” task is from one of this work’s authors (HMS). Panel **b)** depicts scan collection for the National Consortium on Alcohol and NeuroDevelopment in Adolescence (NCANDA) study. A structural and resting-state functional scan were completed at baseline and five years of annual follow-up visits. Panel **c)** depicts scan collection for the Brain Networks and Mobility (BNET) study. A structural and restingstate functional scan were completed at baseline and an 18-month follow-up visit.

The first dataset used in the present work was the Midnight Scan Club (MSC) study (Gordon et al., 2017). MSC includes n=10 (5 female) right-handed, young adult participants. The average age of the sample was 29.1 (standard deviation = 3.3) years, and participants had completed 20.7 (standard deviation = 3.0) years of education on average.

All MRI scans in MSC were collected over twelve sessions on separate days, with each session beginning at midnight. All scans were performed with a Siemens TRIO 3T MRI scanner. Structural MRI images were collected over the course of two days. The full structural imaging protocol is described in detail in the original study. Briefly, four T1-weighted images (sagittal, 224 slices, 0.8 mm isotropic resolution, TE = 3.74ms, TR = 2400ms, TI = 1000ms, flip angle = 8 degrees) were collected for each participant, two per structural scanning day. The only structural images used in the present work were the first T1-weighted images collected on the first day of structural scanning for each participant.

Ten functional scanning sessions were completed on separate days. Each full scanning session lasted approximately 1.5 hours and included resting-state and several tasks. In the present work, only the resting-state, “Faces” task, and “Words” task were used. The Faces task involved distinguishing between male and female faces as they are presented. The Words task consisted of judging whether a presented word was concrete (e.g., “puppy”) or abstract (e.g., “love”). Each fMRI session began with thirty minutes of resting-state, with participants being instructed to fixate on a white crosshair presented on a black background for the full thirty minutes. The remainder of the scan was spent completing various tasks, including Faces and Words. All functional imaging used a gradient-echo EPI sequence (TR = 2200ms, TE = 27ms, voxel dimensions = 4mm x 4mm x 4mm, 36 slices). A gradient echo field map sequence was acquired for each session with the same scan parameters as the functional image sequence.

The second dataset used in this work was from the National Consortium on Alcohol and NeuroDevelopment in Adolescence (NCANDA) study (Brown et al., 2015). NCANDA is a longitudinal multi-site study of n=831 participants aged 12-21 years old at baseline. It is included in analyses in the present work because it represents a dataset that has a large sample size that is comparable to prior work investigating fingerprinting in clustered brain “states” (Cutts et al., 2023; Liu et al., 2018; Van De Ville et al., 2021). It also has many scanning visits per participant and has previously been used for analyses of brain network dynamics (McIntyre, Khodaei, et al., 2025). The present work included only participants from the largest study site (UC San Diego) to avoid confounds from known scanner effects in the NCANDA study (Muller-Oehring et al., 2018; Zhao et al., 2021). Participants completed structural and resting-state functional MRI scans at annual visits throughout the study. In the present work, scans from baseline through follow up year 5 were included in analyses. While NCANDA has high year-to-year retention (Nooner et al., 2020), some participants have unequal numbers of functional scans due to missed visits or study dropout. This work included only participants with available structural and functional scan data for baseline through year 5 follow-up. These inclusion criteria (being enrolled at the UC San Diego site and having six consecutive years of scan data) yielded n=100 (56 female) participants from the NCANDA study aged 15.4 years on average at baseline (standard deviation = 1.8).

The UC San Diego site used a 3T GE Discovery MR750 scanner. High resolution (0.9375mm x 0.9375mm x 1.2mm) T1-weighted images were acquired using an Inversion Recovery-Spoiled Gradient Recalled (IR-SPGR) echo sequence (TR = 5.912ms, TE = 1.932ms, 146 slices, acquisition time = 7m14s). Resting-state BOLD-weighted images were collected using a gradient-recalled EPI sequence (TR = 2200ms, TE = 30ms, voxel dimensions = 4mm x 4mm x 5mm, 32 slices, acquisition time = 10m03s).

The third dataset used in this work was from the Brain Networks and Mobility (BNET) study (Laurienti et al., 2023; McIntyre, O’Donnell, et al., 2025). BNET was a longitudinal study of community-dwelling, cognitively normal older adults aged 70 and older. Participants underwent structural and resting-state functional MRI scanning at baseline and at a 30-month follow-up visit. Of the 192 participants included at baseline, n=173 participants returned for the follow-up scanning visit. BNET is included in analyses for the present work because it has more participants than MSC (n=10) or NCANDA (n=100). It is also comprised entirely of older adults and therefore will assess a fingerprinting feasibility in a different demographic than the prior two studies. For both baseline and follow-up scanning visits, all brain images were collected on a Siemens 3T Skyra MRI scanner equipped with a 32-channel head coil. Each scan session lasted approximately one hour. An anatomical image was collected using T1-weighted 3D volumetric MPRAGE sequence (TR = 2000ms, TE = 2.98 ms, number of slices = 192, 1.0mm isotropic voxels, FOV = 256mm, scan duration = 312s). Resting-state functional MRI data were collected using a blood oxygenation level-dependent (BOLD) (Ogawa et al., 1990) weighted echo planar imaging (EPI) sequence (TR = 2000ms, TE = 25ms, number of slices = 35, voxel dimension = 4mm x4mm x 5mm, FOV = 256mm, duration = 7m14s).

MSC was approved by the Washington University School of Medicine Human Studies Committee and Institutional Review Board. NCANDA was approved by the Institutional Review Boards of the five study sites (UC San Diego, SRI International, Oregon Health and Sciences University, Duke University, and the University of Pittsburgh Medical Center). All BNET participants gave written informed consent to participate in all study activities. BNET was approved by the Institutional Review Board (IRB) of the Wake Forest University School of Medicine (IRB protocol #IRB00046460; approval date: August 27, 2020).

### 2.2 Image Preprocessing

All image preprocessing was completed in SPM 12 (https://www.fil.ion.ucl.ac.uk/spm/) except where specifically noted otherwise. For both each of the three studies, preprocessing began with unified segmentation (Ashburner & Friston, 2005) of T1-weighted structural images using standard six tissue priors while simultaneously warping images to Montreal Neurological Institute (MNI) standard space. We note that while MSC and NCANDA include relatively young participants whose brains tend to warp well to the MNI template using SPM, BNET includes older participants whose brains often do not have satisfactory warps when SPM is used.

Therefore, BNET differed from MSC and NCANDA in the method used to warp images to MNI standard space. For BNET participants, structural images were masked, visually inspected and manually cleaned to remove any remaining non-parenchymal tissues using MRIcron software (https://www.nitrc.org/projects/mricron). Two observers manually checked masked images to ensure accurate full-brain coverage. The masked, cleaned T1-weighted images were spatially normalized to the MNI template using Advanced Normalization Tools (ANTs, https://antsx.github.io/ANTs/).

The first 5 volumes (11.0 s) of functional scans for NCANDA and MSC and the first 10 volumes (20.0 s) of functional scans for BNET were removed to allow signal to reach equilibrium. The remaining volumes (813 for MSC rest, 116 each for MSC Faces and Words tasks, 269 for NCANDA, and 207 for BNET) were corrected for slice-time differences and *B*_0_ distortion, then realigned to the first volume. Slice-timing correction was completed within SPM with the first slice collected in each volume as the reference slice. Functional images were registered to each participant’s T1 image, then warped to MNI space. To remove physiological noise and low frequency drift, functional data was filtered using a band-pass filter (0.009 – 0.08 Hz). Average whole brain gray matter, white matter, and cerebrospinal fluid signals, along with the six degrees of freedom motion parameters obtained from realignment, were regressed from the functional data. To further account for head motion, volumes with motion (at least 0.5mm of framewise displacement) coupled with BOLD signal change (at least 5% change in BOLD from the previous frame) were identified and represented as a binary vector of affected brain volumes (Power et al., 2014). This vector was then used as an additional motion regressor. This motion correction approach allowed retention of the entirety of each participant’s functional scan, which was essential for generating dynamic networks. We note that in the MSC study, participant MSC08 experienced a high degree of drowsiness during the scan and had relatively high levels of motion.

Functional images were then parcellated to the Schaefer 100, 200, 500, and 1000-node 17-subnetwork atlases (Schaefer et al., 2018). Thus, each functional scan was represented in four different parcellation schemes. Reslicing the 1mm x 1mm x 1mm 1000-node Schaefer atlas to the voxel dimensions of the functional scans in the MSC and NCANDA datasets resulted in nodes without any voxels (see Supplementary Figure 1). Therefore, the 4mm x 4mm x 4mm reslicing of the Schaefer 1000-node atlas (for MSC) included only 999 nodes. The 4mm x 4mm x 5mm reslicing of the Schaefer 1000-node atlas (for NCANDA and BNET) included only 998 nodes. For simplicity, both resliced versions of the atlas are still referred to as the Schaefer 1000-node atlas. The number of voxels per node in each parcellation of the resliced atlases are shown in Supplementary Figure 2.

### 2.3 Static and Dynamic Functional Networks

Figure 2 demonstrates how dynamic networks were generated and compares temporally collapsed (by summing across the time dimension) dynamic functional networks to traditional static networks generated using pairwise Pearson correlation of the BOLD time series for each pairing of *N* nodes. This yielded an *N*x*N* functional connectivity matrix for each atlas parcellation of each functional scan.

**Figure 2.**
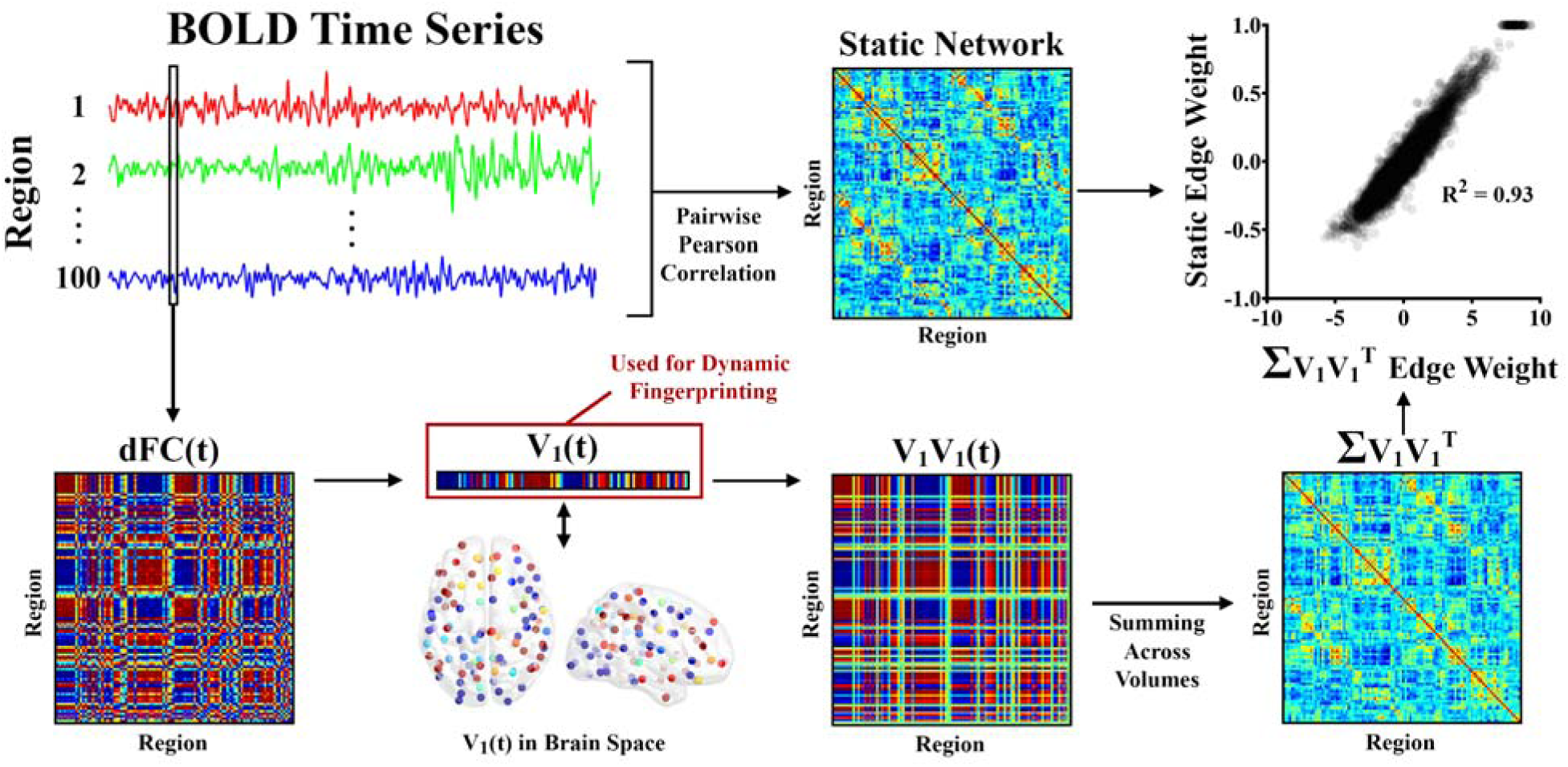
Dynamic networks were generated by finding the phase coherence between all nodes, resulting in a *dFC* matrix for each volume *t*. The leading Eigenvector of *dFC(t), V1(t)*, was identified for each volume *t* and used in dynamic fingerprinting. *dFC(t)* can be approximated by finding the outer product of *V1(t)*, *V1V1(t)*. Summing the edge values across all volumes of *V1V1* results in a matrix with edges that are almost perfectly correlated with the static network generated for the same time series data using Pearson correlation. In all matrices, warm colors indicate positive connections while cool colors indicate negative connections.

To generate dynamic functional networks, BOLD Phase Coherence Connectivity (Deco et al., 2017; Deco & Kringelbach, 2016; Glerean et al., 2012; Ponce-Alvarez et al., 2015) was used on each functional scan to generate a dynamic functional connectivity tensor (dFC). The dFCs are of size *N*x*N*x*T*, where *N* is the number of nodes and *T* is the number of volumes in the scan. To generate phase coherence matrices, the BOLD phase θ*(i,t)* of each node *i* at each volume *t* is first estimated using the Hilbert transform. The Hilbert transform is known to introduce an edge artifact at the beginning and end of time series. To avoid confounds from these artifacts, the first and last five volumes of each scan were removed after applying the Hilbert transform. This resulted in 803 volumes remaining in MSC resting-state networks, 106 volumes for MSC task networks, and 259 volumes remaining in NCANDA networks. The phase coherence between nodes *i* and *j* at volume *t* is obtained according to:

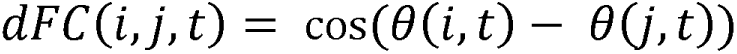

From this equation, it follows that if nodes *i* and *j* have temporally aligned BOLD signals at volume *t*, the difference between their phases will be near 0, and dFC(*i,j,t*) will be approximately 1 (because cos(0) = 1). On the other hand, if *i* and *j* are orthogonal (i.e., if their phases differ by 90 degrees), dFC(*i,j,t*) will be 0 (because cos(90) = 0).

Each dFC volume has *N*(N-1)/2* unique edges – therefore, dFCs are extremely high dimensional, especially in brain parcellations with many nodes. As a result, extensive analyses with the dFC tensors themselves are not computationally feasible. Therefore, in this work dynamic connectivity is represented by the leading eigenvector *V_1_(t)* of the dFC of each volume (Cabral et al., 2017). By focusing on the leading eigenvector of each volume, the dimensionality of the volume is reduced from *N*(N-1)/2* to *N* while still capturing at least 50% of the variance from the dFC. The dFC of each volume is approximately reconstructed by the outer product of the leading eigenvector, *V_1_V_1_(t)*. Collapsing *V_1_V_1_*across time (by summing across volumes for each cell) results in a matrix that is almost perfectly correlated with the scan’s static functional network.

In summary, phase coherence allows estimation of connectivity at every individual volume of a scan. The estimates from phase coherence almost perfectly recreate information present in traditional static networks. The difference is that phase coherence provides an estimate for the connectivity at individual volumes, whereas static networks based on Pearson correlation requires many volumes for their generation and provide an estimate of the average connectivity over all volumes.

### 2.4 Fingerprinting – MSC Rest

Prior work established the feasibility of brain “fingerprinting” using static networks (Finn et al., 2015). In that work, each participant had two static networks, each generated from fMRI time series from separate days. One day of scans was used to make a pool of “database” networks with each participant having one scan in this pool. The other day of scans made a separate pool of “target” networks, again with each participant having one scan in the target pool. To demonstrate the fingerprinting principle, the investigators identified the database network that was most similar (quantified using Pearson correlation) to each target network. The owner of the most similar database network was predicted to be the owner of the target network. Using this approach, the identity of the target network’s owner was identified with >90% accuracy.

In the present work, the fingerprinting approach is extended to dynamic networks. Specifically, this work evaluates the extent to which a single volume of a dynamic network – generated with phase coherence - can identify an individual. The first analysis used the resting-state scans from the MSC dataset. Figure 3 illustrates how the dynamic fingerprinting approach was used in this scenario.

**Figure 3.**
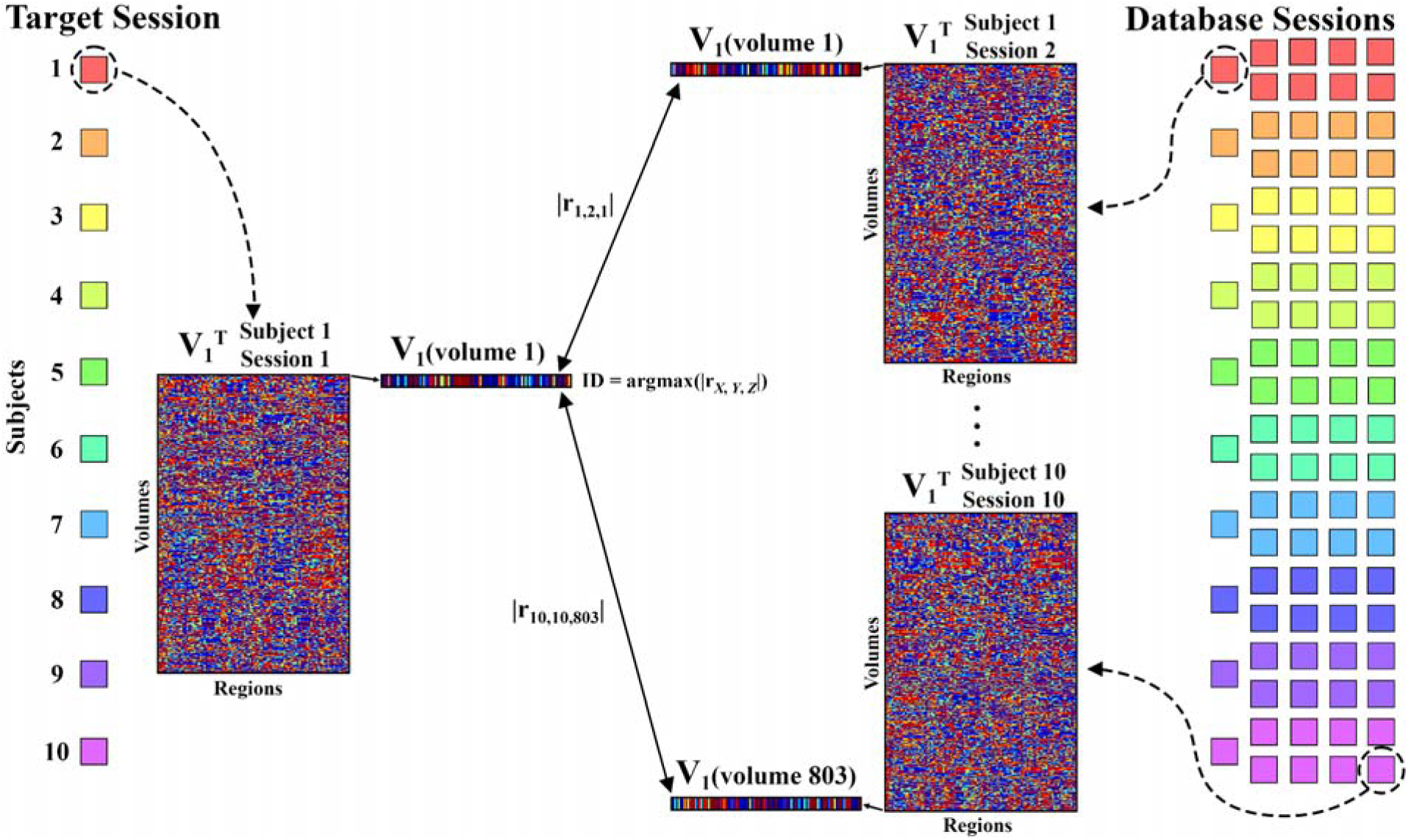
This schematic demonstrates how subject identity was identified for volume 1 of session 1 from Midnight Scan Club subject 1. Each target and database scan (squares with unique colors for participants) was represented by V_1_, a matrix of the leading eigenvector for each volume of the scan. Each volume of the target scan V_1_ was compared to each volume of V_1_ for each database scan by finding the absolute value of the Pearson correlation coefficient |r_x,y,z_| where *x* is the database participant identity, *y* is the database scan session, and *z* is the volume of the database scan. The database volume with the most similar leading eigenvector to each target scan volume was identified (argmax()). The owner of the most-similar database scan volume was predicted to be the owner of the target volume. This process was repeated for every possible target volume.

A single session was designated to make a target pool (one scan per participant). Remaining scans (nine per participant) acted as a database pool. The similarity between each target volume and each database volume was quantified by calculating the absolute value of the Pearson correlation between their leading eigenvectors. The absolute value of the Pearson correlation was used because two eigenvectors with Pearson’s *r* equal to -1 yield identical *V_1_V_1_*matrices – therefore, high |*r*| values indicate similar phase coherence patterns between two volumes. For each target volume, the most similar database volume was identified. The owner of the most similar database volume was predicted to be the owner of the target volume. For each target scan, the number of target volumes that correctly identified the owner of the target scan was recorded. Every possible combination of target-database sessions was tested such that every scan was considered as the target scan once.

Identification accuracy was quantified at two levels. The first level evaluated identifications from individual volumes. At this level, accuracy was quantified as the percentage of target scan volumes that correctly identified the owner. The second level was for entire scans. At this level, the participant with the most volumes suggesting that they were the owner of the scan was the predicted owner of the target scan. For this level, a single accuracy measure was identified across the sample by calculating the percentage of target scans that correctly identified the participant.

To determine whether single-volume identification accuracy was statistically significant, a linear mixed effects model was fit to test the null hypothesis that identification accuracy was due to random chance. In this model, each observation represented a different target scan, resulting in 100 observations (10 participants x 10 sessions). The outcome in this model was the number of target volumes that correctly identified their owner. The identities of target scan owners were included as random effects to account for repeated subjects in the analysis. We assessed whether the number of correct volumes that would be identified by random chance (803 volumes per target scan / 10 possible participant labels = 80.3 volumes) was outside of the 95% confidence interval for the linear mixed effects model’s intercept. All linear mixed effects models in this work were modeled using the MATLAB (version 2025a) function ‘fitlme’ – scripts used in analyses are available on github (see *Data and Code Availability* below).

Additionally, significance for the single-volume and full-scan levels were assessed with nonparametric permutation testing. For 1000 permutations, the identity labels of database scans were randomized, and identification accuracy for the two levels using the permuted database labels was calculated. *p*-values were calculated by dividing the count of permutations with identification performance equal or superior to the actual identification performance divided by the total number of permutations.

### 2.5 Fingerprinting – NCANDA Rest

To assess fingerprinting feasibility using phase coherence measures in a second, independent sample, the full process described in the previous section was repeated in the NCANDA sample. In addition to providing an independent sample, NCANDA follows participants through several years of adolescence and young adulthood, meaning that this sample provides a valuable perspective on the robustness of single-frame connectivity identification longitudinally during adolescent neurodevelopment.

Fingerprinting for the NCANDA sample followed the same process as MSC with minor adjustments. While MSC had ten sessions per subject, NCANDA had six sessions per subject. Therefore, each NCANDA target scan was compared against five database scans as opposed to the nine database scans used in MSC. Additionally, while MSC had n=10 participants with 803 volumes per scan, NCANDA had n=100 participants with 259 volumes per scan. Therefore, by random chance, only 2.59 volumes (259 volumes per scan / 100 participants) were expected to correctly identify the participant on average. To determine statistical significance, we evaluated whether 2.59 was outside the 95% confidence intervals for the linear mixed effects model intercepts. The same nonparametric permutation testing approach from the MSC analyses was also applied in NCANDA.

### 2.6 Fingerprinting – BNET Rest

To assess fingerprinting feasibility using phase coherence measures in a third, larger independent sample, the full process described in the previous sections was repeated for BNET. Fingerprinting for the BNET sample followed the same process as MSC and NCANDA, but because BNET only had two scans, the effect of different numbers of database scans was not tested. Additionally, BNET had n=173 participants with 197 volumes per scan. Therefore, by random chance, only 1.14 volumes (197 volumes per scan / 173 participants) were expected to correctly identify the participant on average. To determine statistical significance, we evaluated whether 1.14 was outside the 95% confidence intervals for the linear mixed effects model intercepts. The same nonparametric permutation testing approach from the MSC and NCANDA analyses was also applied in BNET.

### 2.7 Fingerprinting – Static Networks

For each of the three studies included in this work, the fingerprinting performance of the resting-state static networks is assessed in supplemental analyses for comparison to the performance yielded by individual volumes and full scans using the dynamic fingerprinting approach. Static fingerprinting analyses were carried out according to the protocol established in prior work (Finn et al., 2015).

### 2.8 Fingerprinting – Atlas Parcellation

To determine how the size of the network impacts identification accuracy, the full fingerprinting process described above was attempted with four different parcellations of the Schaefer atlas for MSC and NCANDA (see *Image Preprocessing*). Linear mixed effects models were fit for MSC and NCANDA separately. The outcome variable in models were the number of volumes correctly matched to a participant for a given target scan. Atlas approach was a categorical fixed effect, and subject identity was included as a random effect to account for repeated measures.

Additionally, to determine whether differences in identification performance between the highest (1000 node) and lowest (100 node) resolution atlases were due to the number of nodes or the size of nodes, we assessed identification performance in the MSC dataset using only 100 nodes from the Schaefer 1000 atlas. With this approach, performance of 100 high-resolution nodes (from the Schaefer 1000 atlas) could be compared against 100 low-resolution nodes (from the Schaefer 100 atlas). With the number of nodes being constant between the two approaches, performance differences should either be due to differences in brain coverage or differences in node size. For 100 iterations, 100 nodes were randomly sampled from the Schaefer 1000-node atlas. The number of nodes representing each of the 17 Yeo atlas subnetworks (Yeo et al., 2011) was held consistent with the original Schaefer 100-node atlas. Full fingerprinting analyses were carried out with the dynamic networks resulting from each of the down sampled 100-node atlases. To determine whether the down sampled 100-node atlases differed in performance from either the original 100 or 1000 node atlases, a linear mixed effects model was fit with the atlas approach (100 nodes, 1000 nodes, or 100 nodes down sampled from 1000 nodes) as a categorical fixed effect and target subject identity as a random effect.

We note that averaging time series across many voxels (as in a large atlas region) often has the effect of smoothing BOLD signal, which reduces the “noise” in the signal and enhances temporal signal to noise ratio (tSNR). In supplemental analyses, we assess differences in the average temporal signal to noise ratio (tSNR) in nodes from the 100- versus 1000-region Schaefer atlas.

### 2.9 Fingerprinting – Number of Database Scans

To determine how the number of database sessions affects identification accuracy, the full fingerprinting process described above was carried out with different numbers of database scans. For both MSC and NCANDA, accuracy when only one database scan was available was assessed. Each possible combination of target-database scans was evaluated. This means that for each subject, there were 90 iterations of target-database scans in MSC and 30 iterations of target-database scans in NCANDA. Additionally, performance was tested for every possible combination of five database scans in MSC and three database scans in NCANDA. The number of possible target-database scan combinations is equal to *n* choose *k*, where *n* is the number of possible database scans (9 for MSC, 5 for NCANDA) and *k* is the number of chosen database scans (5 for MSC, 3 for NCANDA). Therefore, for each subject, there were 126 iterations of target-database scans in MSC and 10 iterations of target-database scans for NCANDA.

Linear mixed effects models were used in similar fashion to the linear mixed effects models in the atlas parcellation analyses described above. Instead of atlas being a fixed effect, the number of database scans for each target scan was now the categorical fixed effect.

### 2.10 Fingerprinting – Gray Matter Signal Regression

Global brain signal regression is a highly debated topic among researchers working with fMRI data (Liu et al., 2017). In this work, we elected to remove global signal by regressing average whole brain gray matter signal from voxel time series. However, because many individuals elect not to carry out this preprocessing step, we also replicated dynamic fingerprinting analyses in MSC using time series data that was preprocessed in the same way but omitted the whole-brain average gray matter signal regression step. Analyses were recreated for the Schaefer 100 and 1000 atlas parcellations.

### 2.11 Fingerprinting – Motion Artifact Effect

It is common practice to remove volumes that have high motion from time series prior to generating correlation-based networks due to artifacts that can be introduced due to motion (Power et al., 2014). Because this study was focused on brain network dynamics, every timepoint was retained for analyses despite some volumes having high motion. While several steps were applied in preprocessing to address motion artifacts, ultimately the signal from these volumes may still differ from the signal of volumes with low head motion. To assess the effect of head motion on fingerprinting performance, we identified volumes that would have been removed in the motion scrubbing procedure (see *Image Preprocessing* for more detail). For each of the three studies individually, we identified the identification accuracy for these high-motion volumes as compared to identification accuracy for volumes that would have been retained in the motion-scrubbing procedure.

### 2.12 Fingerprinting – Task Identification

Determining whether the fingerprinting approach could distinguish between tasks was also of interest. Using the atlas parcellation with the best participant identification performance and nine database sessions of three tasks (Faces, Words, and Rest) each, the accuracy for task identification of a target scan was assessed. Both the Faces and Words networks featured 106 volumes each. While the resting-state scans were substantially longer, only the first 106 volumes of the resting-state scan for each visit were retained for this analysis to ensure that each task had an equal number of volumes per scan.

Figure 4 demonstrates how the dynamic fingerprinting approach was altered to identify tasks. Task identification was first assessed within participant (Figure 4a). One of the ten sessions was designated the target scan with the nine other sessions being the database – all target and database scans belonged to the same participant, and all three tasks were present in the database. The fingerprinting process was followed, but the label of interest was now the task of the target scan rather than the identity of the participant. The number of target volumes with correctly identified tasks were counted. Every combination of task and target-database sessions was used for every participant. A linear mixed effects model was then fit separately for each task, with each target session for the task being included as an observation. The outcome of each observation was the number of volumes with correct task identification. Subject identity was included as a random effect to account for repeated measures. To assess statistical significance, the number of task classification expected to be correct by random chance (106 volumes / 3 tasks = 35.3 volumes) was compared against the 95% confidence interval for the intercept of the linear mixed effects model.

**Figure 4.**
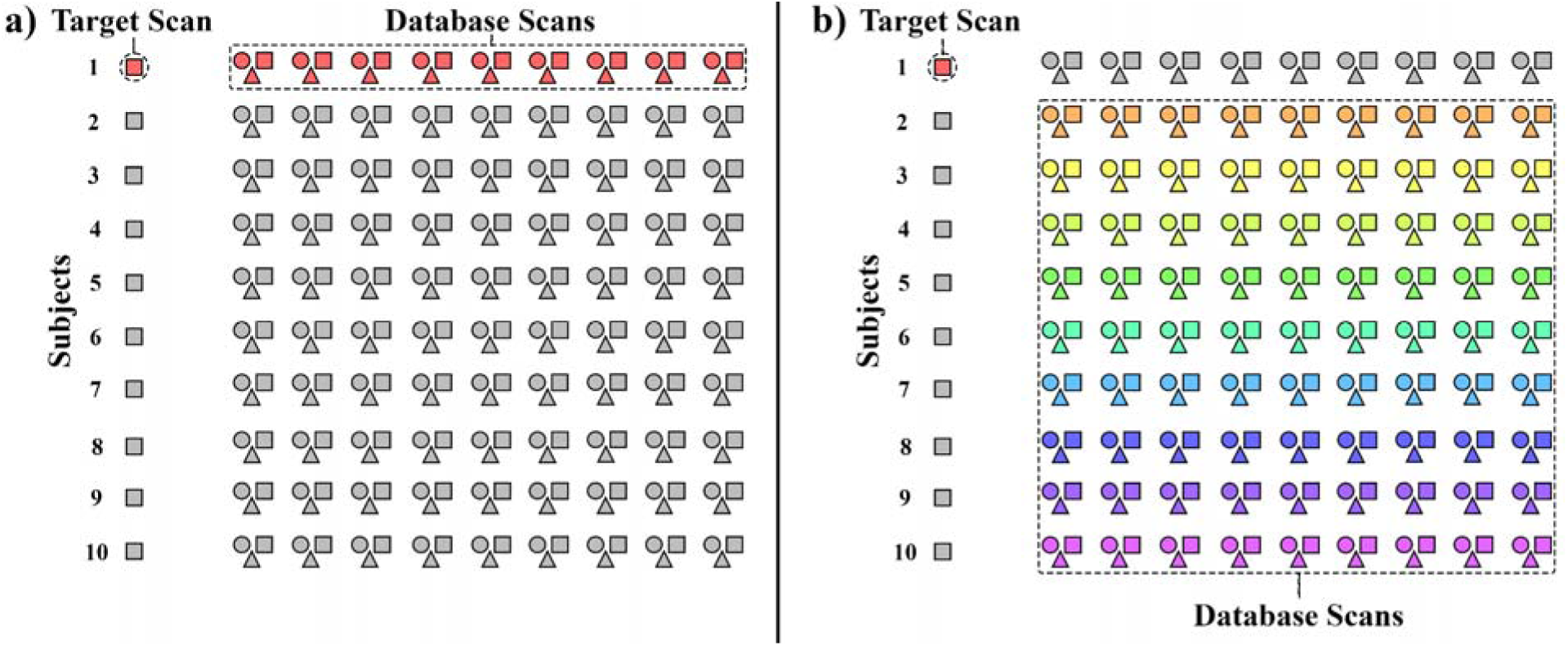
This schematic demonstrates how database scan pooling was modified for task identification. In this example, each target scan is from the participant at rest. Each participant has scans from other sessions for rest (squares), Faces task (triangles) and Words task (circles). a) First, only scans from the same participant as the target scan were included in the database pool. The database volume that was most similar to each target volume was identified. The task that was being completed in the database volume was the predicted task for the target volume. b) Next, only scans from non-target participants were included in the database pool. The same task classification process was repeated with that pool.

Separately, task identification was assessed between participants (Figure 4b). This analysis followed the same approach as within-subject task identification, but the database scans were now from the other nine participants of MSC (i.e., not the same participant that the target scan came from). Non-target sessions from the owner of the target scan were excluded from the database. As in the within-subject analysis, identification of all tasks as targets were evaluated in separate linear mixed effects models.

Finally, the performance of within versus between-subject task identification was compared. The motivation of this analysis was to assess the extent to which single-volume connectivity patterns occupied during distinct cognitive tasks are consistent across a sample or unique to individuals. A linear mixed effects model was fit separately for each task. In these models, there was an observation for each target session of the within and between-subject approach (200 observations total). For each observation, the outcome was the number of target scan volumes with correctly identified task. A binary marker indicating whether the observation followed the within-subject or between-subject approach was included as a fixed effect, and subject identities were included as random effects.

## 3. Results

### 3.1 Participant Identification – MSC

Figure 5 shows bar plots depicting the number of volumes from each participant’s target scans that were predicted to belong to each of the ten MSC participants. Results from the 100-and 1000-node atlas with 9 database sessions are shown. For each target scan, more volumes were predicted to belong to the owner of the target scan than any other subject. Therefore, the full-scan identification accuracy for these participants was 100% (Figure 6 and Supplementary Table 2).

**Figure 5.**
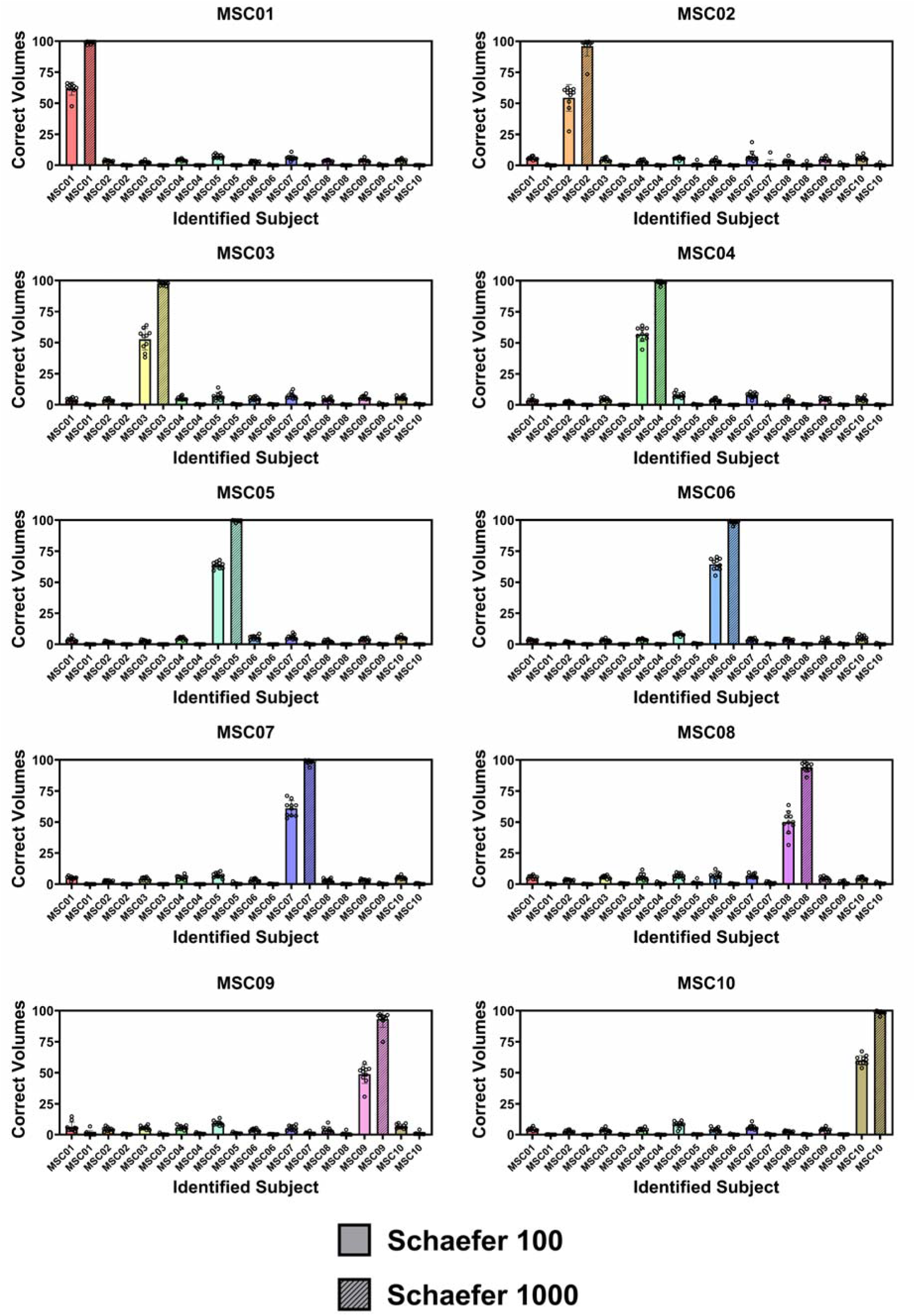
Each bar plot corresponds to a different target participant. Plots show the percentage of total scan volumes predicted to belong to each of the database participants (x-axis) for fingerprinting analyses with the Schaefer 100-node atlas (solid bars) and the Schaefer 1000-node atlas (striped bars). Each bar plot includes data from ten target scans with each dot of each bar representing a different target scan. For every target scan shown above, the participant that was predicted most often was the participant that the target scan belonged to (i.e., the correct participant was always identified when all scan volumes were considered). The accuracy of individual volume identification improved substantially by increasing the atlas parcellation from 100 to 1000 regions.

**Figure 6.**
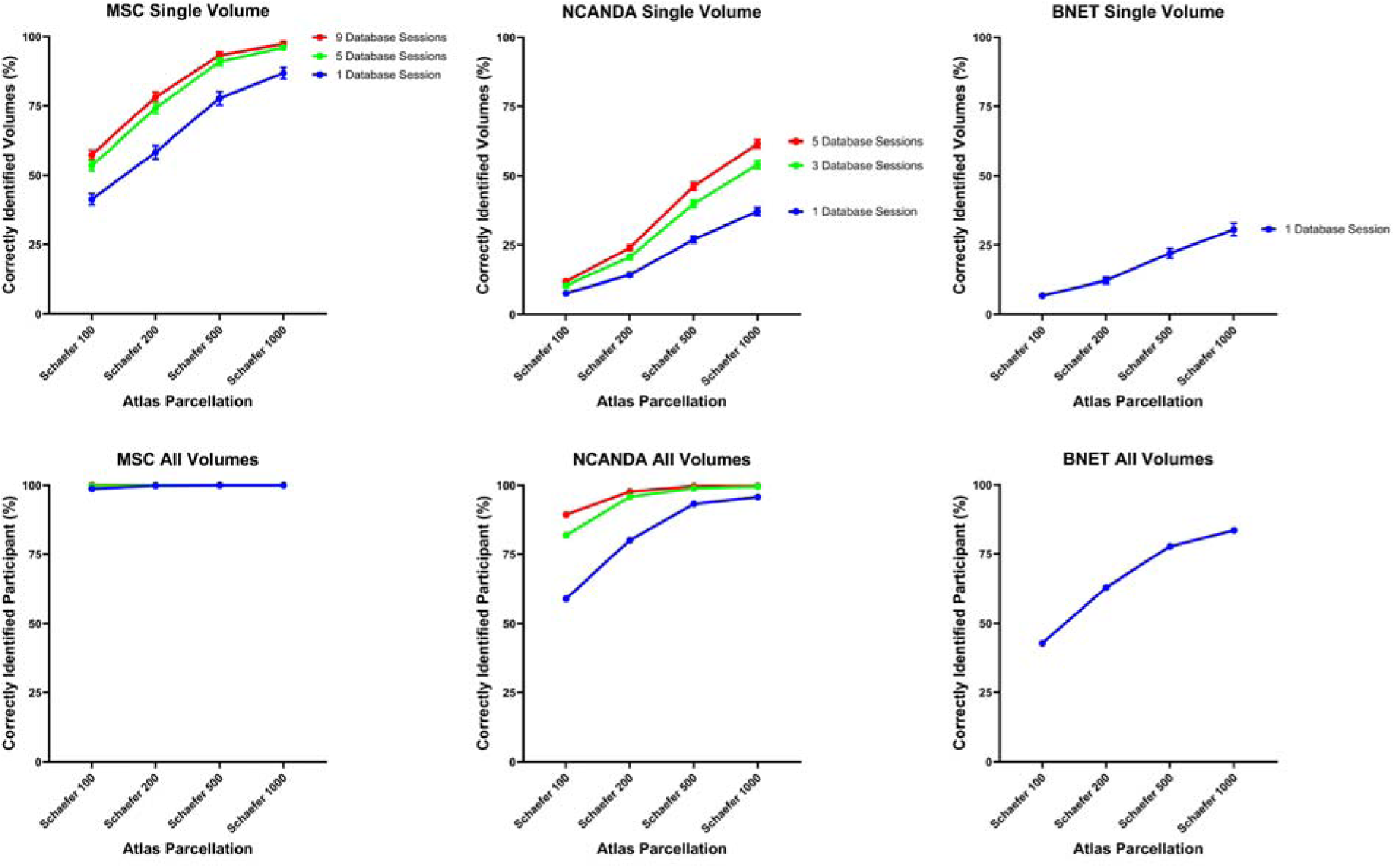
Identification accuracy across the three different studies (MSC, NCANDA, and BNET) with different atlas parcellations (shown on x-axes) and amounts of database scans (shown as different line colors). In the top row, y-axes show the identification accuracy of individual volumes for each study. Error bars represent 95% confidence intervals. In the bottom row, y-axes show the percentage of scans for which the identity of the participant was correctly classified when all volumes from the scan were considered.

Figure 6 and Supplementary Tables 1 and 2 demonstrate the participant identification accuracy for the MSC sample when using individual volumes and full scans. For every combination of atlas parcellation and number of database scans, the number of volumes expected to be correct by chance (80.3) was outside of the 99.9% confidence interval. Therefore, subject identification for every combination was statistically significant with *p* < 0.001 when assessed with this approach. Statistical significance of participant identification was also assessed using permutation testing. Out of all permutations (1000 per combination of atlas and number of database scans), identification accuracy was never greater than or equal to the experimental identification accuracy for the single-volume or full-scan level. Therefore, this approach also showed that participant identification was statistically significant with *p* < 0.001 for every atlas and number of database combination.

While every combination of atlas parcellation and number of database scans had identification performance that exceeded random chance, identification accuracy clearly improved with higher atlas resolution (more nodes) and with more database scans. Linear mixed effects models confirmed that each step in increased resolution (Schaefer 100 → Schaefer 200 → Schaefer 500 → Schaefer 1000) had significantly higher identification rates than the lower resolution, with *p* < 0.001 for each step. Linear mixed effects models also confirmed that each step in additional database scans (1 scan → 5 scans → 9 scans) had significantly higher identification rates than fewer database scans, with *p* < 0.001 for each step.

### 3.2 Participant Identification - NCANDA

Figure 6 and Supplementary Tables 3 and 4 demonstrate the participant identification accuracy for the NCANDA sample when using individual volumes and full scans. For every combination of atlas parcellation and number of database scans, the number of volumes expected to be correct by chance (2.59) was outside of the 99.9% confidence interval. Therefore, subject identification for every combination was statistically significant with *p* < 0.001 when assessed with this approach. Statistical significance of participant identification was also assessed using permutation testing. Out of all permutations (1000 per combination of atlas and number of database scans), identification accuracy was never greater than or equal to the experimental identification accuracy for the single-volume or full-scan level. Therefore, this approach also showed that participant identification was statistically significant with *p* < 0.001 for every atlas and number of database scan combination.

As with the MSC results, identification accuracy improved with higher atlas resolution and with more database scans. Linear mixed effects models confirmed that each step in increased resolution (Schaefer 100 → Schaefer 200 → Schaefer 500 → Schaefer 1000) had significantly higher identification rates than lower resolution, with *p* < 0.001 for each step. Linear mixed effects models also confirmed that each step in additional database scans (1 scan → 3 scans → 5 scans) had significantly higher identification rates than fewer database scans, with *p* < 0.001 for each step.

### 3.3 Participant Identification - BNET

Figure 6 and Supplementary Tables 5 and 6 demonstrate the participant identification accuracy for the BNET sample when using individual volumes and full scans. For every combination of atlas parcellation and number of database scans, the number of volumes expected to be correct by chance (1.14) was outside of the 99.9% confidence interval. Therefore, subject identification for every combination was statistically better than chance with *p* < 0.001 when assessed with this approach. Statistical significance of participant identification was also assessed using permutation testing. Out of all permutations (1000 per combination of atlas and number of database scans), identification accuracy was never greater than or equal to the experimental identification accuracy for the single-volume or full-scan level. Therefore, this approach also showed that participant identification was statistically better than chance with *p* < 0.001 for every atlas.

As with the MSC and NCANDA results, identification accuracy improved with higher atlas resolution. Linear mixed effects models confirmed that each step in increased resolution (Schaefer 100 → Schaefer 200 → Schaefer 500 → Schaefer 1000) had significantly higher identification rates than lower resolution, with *p* < 0.001 for each step.

### 3.3 Atlas Parcellation Down Sampling

The single-volume participant identification accuracy of the 100 iterations of down sampled 100-node Schaefer atlases was 70.0% ± 1.8%. A linear mixed effects model indicated that this performance was superior to the original 100-node Schaefer atlas (Mean accuracy = 57.4%, *p* < 0.001), but inferior to the 1000-node Schaefer atlas (Mean accuracy = 97.4%, *p* < 0.001).

The average tSNR of nodes from the 100- and 1000-node Schaefer atlas was compared for the MSC dataset to assess the extent to which the different parcellations differ in the amount of noise present. Supplementary Figure 3 shows the average nodal tSNR for both atlas parcellations in all 100 MSC scans. The average tSNR for the Schaefer 100 atlas was 576.2 (standard deviation = 113.5) while the average tSNR for the Schaefer 1000 atlas was 396.8 (standard deviation = 70.7). Paired t-tests revealed that this difference was statistically significant (*p* < 0.001).

### 3.4 Supplemental Resting-State Fingerprinting Analyses

Supplementary Tables 7 and 8 show results from repeating the dynamic fingerprinting process in the MSC dataset without regressing average whole brain gray matter signal from time series data. The results were not meaningfully different from the analyses of data that was preprocessed with gray matter regression (Supplementary Tables 1 and 2).

We also assessed whether participant identification was more successful in high-motion or low-motion volumes to assess whether fingerprinting success was driven by head motion in the scanner. Supplementary Table 9 shows the percentage of high-motion versus low-motion volumes for which the participant was correctly identified in the MSC dataset. Supplementary Tables 10 and 11 show the same thing for the NCANDA and BNET studies, respectively. Across all three studies, the identification performance of high-motion volumes was clearly lower than the identification performance of low-motion volumes.

Finally, in order to compare dynamic fingerprinting performance to static fingerprinting performance, we determined the accuracy with which participants could be identified based on their static networks. Results for the MSC, NCANDA, and BNET studies are shown in Supplementary Tables 12-14, respectively. In the MSC dataset, there did not appear to be a meaningful difference in performance between the static network fingerprinting and the dynamic fingerprinting when all volumes were considered. In NCANDA and BNET, a trend emerged such that the static networks appeared to have better fingerprinting performance than the dynamic networks when the Schaefer 100 atlas was used. However, when higher parcellation atlases were used, the superior performance of the static networks was not observed.

### 3.5 Task Identification

Task identification performance is shown in Figure 7 and Supplementary Table 15. Within subjects, the average single-volume task identification accuracy was 58.4% for the Faces task, 46.4% for the Words task, and 60.3% for resting state. Linear mixed effects models showed that identification for all three tasks was superior to random chance (33.3%) with *p* < 0.001. Between subjects, the average single-volume task identification accuracy was 49.9% for the Faces task, 32.1% for the Words task, and 51.4% for resting state. Linear mixed effects models showed that identification rates for Faces and resting state were superior to random chance (33.3%) with *p* < 0.001. However, identification for the Words task was not significantly different from chance performance.

**Figure 7.**
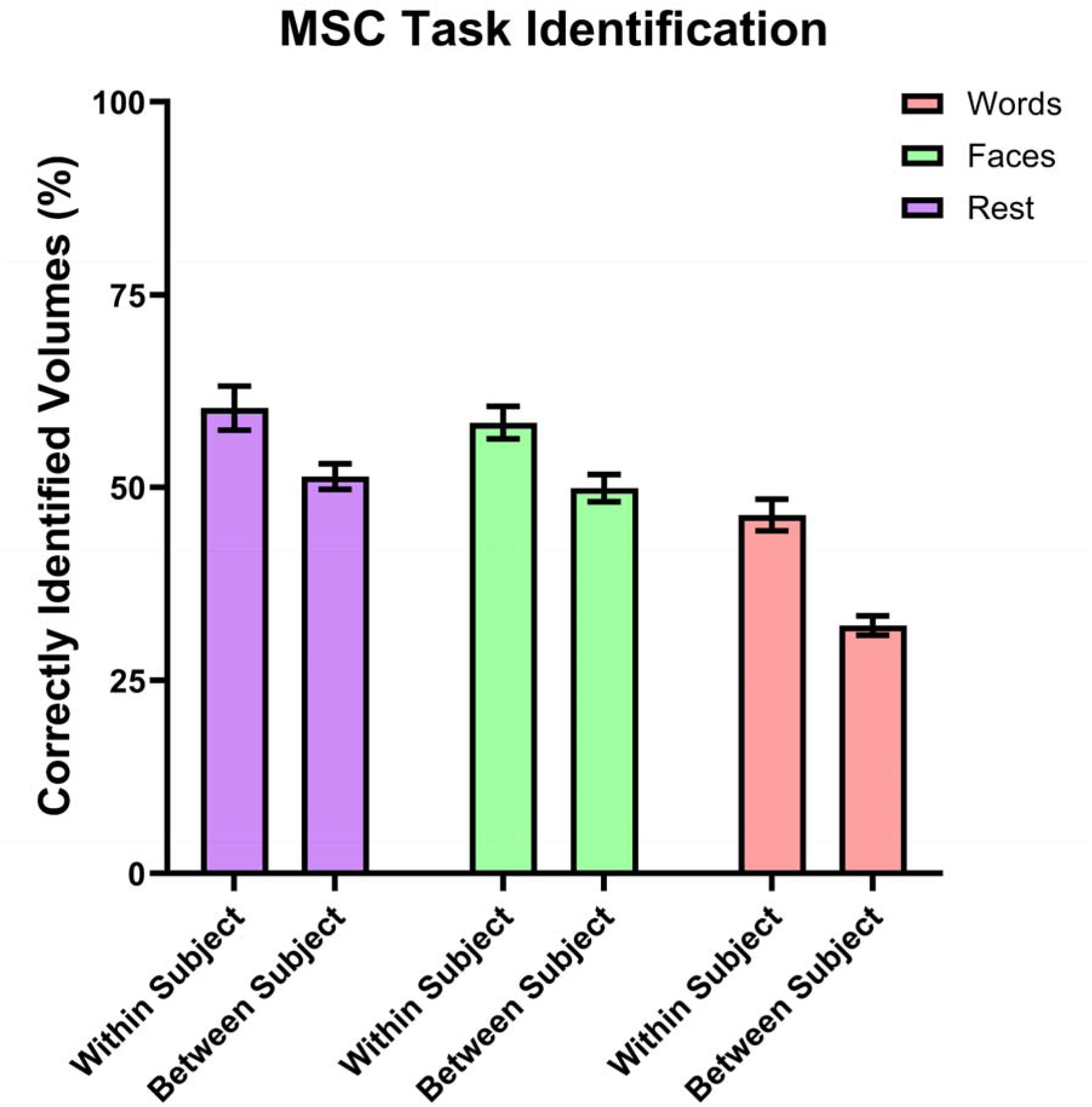
Percentage of volumes for which the task was correctly identified. Error bars represent 95% confidence intervals. By random chance, task identification for individual volumes would be expected to be correct 33.3% of the time. For each of the three tasks, within-subject task identification was statistically better than between-subject task identification.

Additionally, linear mixed effects models assessed whether task identification accuracy within subject was different from task identification accuracy between participants. These models revealed that within-subject task identification was superior to between-subject task identification for Faces, Words, and Rest, with *p* < 0.001 for each task.

## 4. Discussion

The idea that brain function is individualized and ever-changing is not new, nor was it new when functional connectome fingerprinting (Anderson et al., 2011; Finn et al., 2015), precision functional mapping (Gordon et al., 2017), or methods for studying dynamic brain networks (Allen et al., 2014; Chang & Glover, 2010; Handwerker et al., 2012; Hutchison et al., 2013) were first introduced and popularized. Individuality and temporal variability in brain function were considered core features of consciousness (James, 1892) almost a century before the discovery of BOLD signal made the widespread, empirical study of brain physiology in living humans possible (Ogawa et al., 1990). The major contribution of the more recent studies was that they established approaches to quantify individuality and time-variance in brain network models of BOLD signal. Even more recent studies have begun to show that dynamic brain “states” are likely unique to individuals (Cutts et al., 2023; Ghaffari et al., 2025; Liu et al., 2018; Menon & Krishnamurthy, 2019; Van De Ville et al., 2021). The present work leveraged phase coherence to assess the extent to which the findings from these most recent studies extend to single-frame connectivity across three different studies.

Most studies of brain network dynamics define “states” as recurring patterns of connectivity that are identified at a group level, generally resulting in less than ten states (Allen et al., 2014; Cabral et al., 2017; Shappell et al., 2019; Vidaurre et al., 2018). Subsequently, studies examine how individuals differ in their trajectories through the states that were defined at the group level. Using this approach, trajectories through states can differ between individuals, but the states that individuals occupy cannot. Prior work has shown that if subject-specific “states” are identified using clustering algorithms within-subject, then the identity of the subject can be predicted with high accuracy based on their unique set of states (Ghaffari et al., 2025; Menon & Krishnamurthy, 2019).

Importantly, cluster-based definitions of states are limited in that there is no ground truth as to how many states should be used to represent the range of connectivity patterns present in a dataset. Further, the number of states that different participants occupy in a given scan may differ. To avoid issues relating to clustering many timepoints into a small number of brain states, in this work we allow each timepoint to retain its own unique connectivity pattern rather than attributing the connectivity from a cluster of frames to any individual frame. Therefore, in this work, the connectivity at every volume is different from every other volume, though some volumes are more similar than others. Using this approach, we find that in the MSC, single-frame connectivity patterns from one participant tend to be most similar to other frame from that same participant than to frames from other participants. This suggests that at a single-frame temporal resolution, connectivity patterns tend to be unique to individuals.

The general conclusions from MSC resting-state scans extended to the NCANDA and BNET studies. Compared to MSC, single-volume and full-scan identification performance in the NCANDA and BNET studies was lower, which was not surprising considered that these two studies had substantially larger sample sizes and fewer database volumes than MSC. Nevertheless, participant identification at both the single-volume and full-scan level was better than chance in both the NCANDA and BNET studies, which supports that the identity of the participant plays a role in the transient connectivity patterns that we should expect to observe from their BOLD time series.

The NCANDA sample features participants who are adolescents at baseline. Throughout five years of follow-up, these participants are undergoing substantial neurodevelopmental changes in brain structure and function (Bethlehem et al., 2022; Fair et al., 2008; Fair et al., 2009; Fair et al., 2007). Nevertheless, we find that individuals’ single-volume connectivity patterns retain some level of identifiability. Previous work has shown that static networks become distinct as early as infancy (Hu et al., 2022) and that their distinctiveness is reliable across many years in adulthood (St-Onge et al., 2023). The findings from the NCANDA sample in the present work indicate that lifespan connectome fingerprint stability is not limited to static connectomes. Rather, it extends to transient patterns of connectivity. Additionally, the BNET study features participants who are 70+ years old. Identification of individuals in the BNET sample was only slightly less successful than NCANDA when one database scan was used. The slightly lower performance is likely due to BNET having more total participants and fewer volumes per scan to serve as a database. The finding that dynamic fingerprinting in older adults had similar success to adolescents/young adults further supports that dynamic fingerprints may be stable across large portions of the lifespan.

Individual identifiability in resting-state increased with the number of nodes in brain parcellations. A similar result was reported in prior fingerprinting analyses on static networks (Finn et al., 2015). In static networks, increasing the number of nodes in a parcellation has been shown to improve classification of task (Said et al., 2023) and sex (Arslan et al., 2018; Said et al., 2023). We note that supplemental analyses revealed that the time series from regions of the Schaefer 1000 atlas had lower tSNR than regions of the Schaefer 100 atlas, meaning that having smaller nodes resulted in “noisier” signal. However, prior work has also shown that having smaller regions that represent fewer voxels results in higher homogeneity among the voxels in the region (Stanley et al., 2013). Having more homogenous time series from the voxels within a region may indicate that the region is a more meaningful representative of all voxels in the region. Therefore, the choice of parcellation level requires consideration of both signal to noise ratio (which tends to be superior in larger atlas regions) and signal homogeneity (which tends to be superior in smaller atlas regions). We find that while they may be noisier, the dynamic connectivity patterns from smaller atlas nodes were also more unique to individuals than connectivity patterns from larger atlas nodes.

The present work also found that having more scan data in the database scan pool increased individual identifiability. This finding is also in agreement with prior findings from static network fingerprinting (Anderson et al., 2011; Finn et al., 2015). We suggest that smaller amounts of database data may result in lower identification rates due to noise inherent in phase coherence measures (Pedersen et al., 2018) and fMRI data itself. We hypothesize that this effect may be mitigated by increasing the amount of database data. We believe that having more database scans reduces the chances that participants will be misidentified because noise-related similarities in connectivity between different participants should not be consistent across many scan sessions, whereas legitimately similar connectivity patterns should be consistent within subject.

The present work also found that within subject, cognitive task could be identified at a statistically significant rate. Prior studies have used sample-level states to show that the amount of time spent in different states differ in rest versus working memory task (Kurtin et al., 2023) and when interacting with imagery of happy versus unhappy infants (Stark et al., 2020). In the present work, some tasks (resting state and Faces) could be distinguished by comparing volumes across participants. However, identification of each task was superior when comparing volumes within subject. The implication of this finding is that at the single-volume level, the connectivity patterns that support a specific cognitive task may slightly differ by subject.

Task performance was most successful for resting-state. Whereas both the Words and Faces task required participants to be actively engaged and interact with prompts, resting state differed in that it did not require participants to engage with anything besides a crosshair. We interpret the superior task identification performance of resting state scans to be a reflection of this distinction from the other tasks. Because resting state was more different from the other two tasks, it seems to have been easier to identify. This interpretation is supported by prior work that used lower-dimensional embedding to visualize the similarities in activation patterns of individual volumes from participants engaged in various cognitive tasks (Saggar et al., 2018).

The prior study found that while different tasks tended to have very similar activation patterns within subjects, activation during rest tended to be much more unique to rest.

The present work acknowledges several limitations. First, several have suggested that brain network dynamics observed during resting state scans may only reflect signal variability around a static network (Ladwig et al., 2022; Laumann et al., 2017). The goal of this work is not to prove or disprove that the brain should be modeled solely as a dynamic or static network.

Rather, the goal of this work was to determine the extent to which the developing field of dynamic fingerprinting (Cutts et al., 2023; Ghaffari et al., 2025; Liu et al., 2018; Menon & Krishnamurthy, 2019; Van De Ville et al., 2021) might extend to individual volumes in fMRI data. However, given that we have not proven that the brain occupies distinct states that are not explainable by sampling variability, we note that an alternative interpretation to findings in this work may be that the uniqueness of static networks trickles down to the level of connectivity at single volumes estimated using phase coherence.

Another limitation is that while connectivity patterns estimated with phase coherence has better temporal resolution than methods like sliding window correlation, it may also be prone to higher levels of noise (Pedersen et al., 2018). This work cannot answer how the phase coherence measure’s vulnerability to noise may alter individual identification. We hypothesize that identification accuracy would only be diminished by high-frequency noise, as we expect that noise should manifest uniformly across the sample and result in connectivity patterns appearing more similar between individuals. Analyses assessing the identification performance of volumes with high motion indicated that this is likely the case with motion-related noise, but we cannot conclusively say how other sources of physiological noise may impact identification performance. Future work could evaluate this effect in more detail by introducing noise of different distributions to time series prior to attempting fingerprinting.

## Data and Code Availability

Data in this work came from the MSC, NCANDA, and BNET studies. MSC is publicly available here: https://openneuro.org/datasets/ds000224/versions/1.0.4. NCANDA data is available following an application process beginning here: https://ncanda.org/datasharing.php. BNET data is not publicly available as it was collected prior to mandatory public sharing of NIH-funded fMRI data acquisition, and participants therefore did not provide data release agreements. The BNET data can be made available upon request to the authors with appropriate Institutional Review Board approval and data use agreements. MATLAB scripts were written to perform all analyses in this work. These scripts are available on github: https://github.com/ccmcinty/The-individuality-of-single-frame-functional-brain-connectivity-.

## Author Contributions

**Clayton McIntyre:** Conceptualization, Methodology, Software, Formal Analysis, Funding Acquisition, Data Curation, Writing – Original Draft, Writing – Reviewing & Editing, Visualization. **Heather Shappell:** Methodology, Formal Analysis, Funding Acquisition, Writing – Reviewing & Editing. **Mohsen Bahrami:** Methodology, Formal Analysis, Funding Acquisition, Writing – Reviewing & Editing. **Robert Lyday:** Software, Resources, Data Curation, Writing – Reviewing & Editing, Visualization. **Paul Laurienti:** Conceptualization, Supervision, Funding Acquisition, Writing – Reviewing & Editing.

## Funding

Analyses were supported by NIH grants P50AA026117 (PJL), RO1AG052419 (PJL), RO1AG052419-05S1 (PJL), P30AG021332 (PJL), K25EB032903 (HMS), K25AG090707 (MB), and F31AA032409 (CCM), the National Center for Advancing Translational Sciences (IL1TR001420), and the Wake Forest University Translational Science Center

## Declaration of Competing Interests

The authors declare no conflicts of interest.

## Acknowledgements

The authors thank the participants and study teams from the Midnight Scan Club, National Consortium on Alcohol and NeuroDevelopment in Adolescence, and Brain Networks and Mobility studies for providing the data used in this work.

---

**Supplementary Figure 1.**
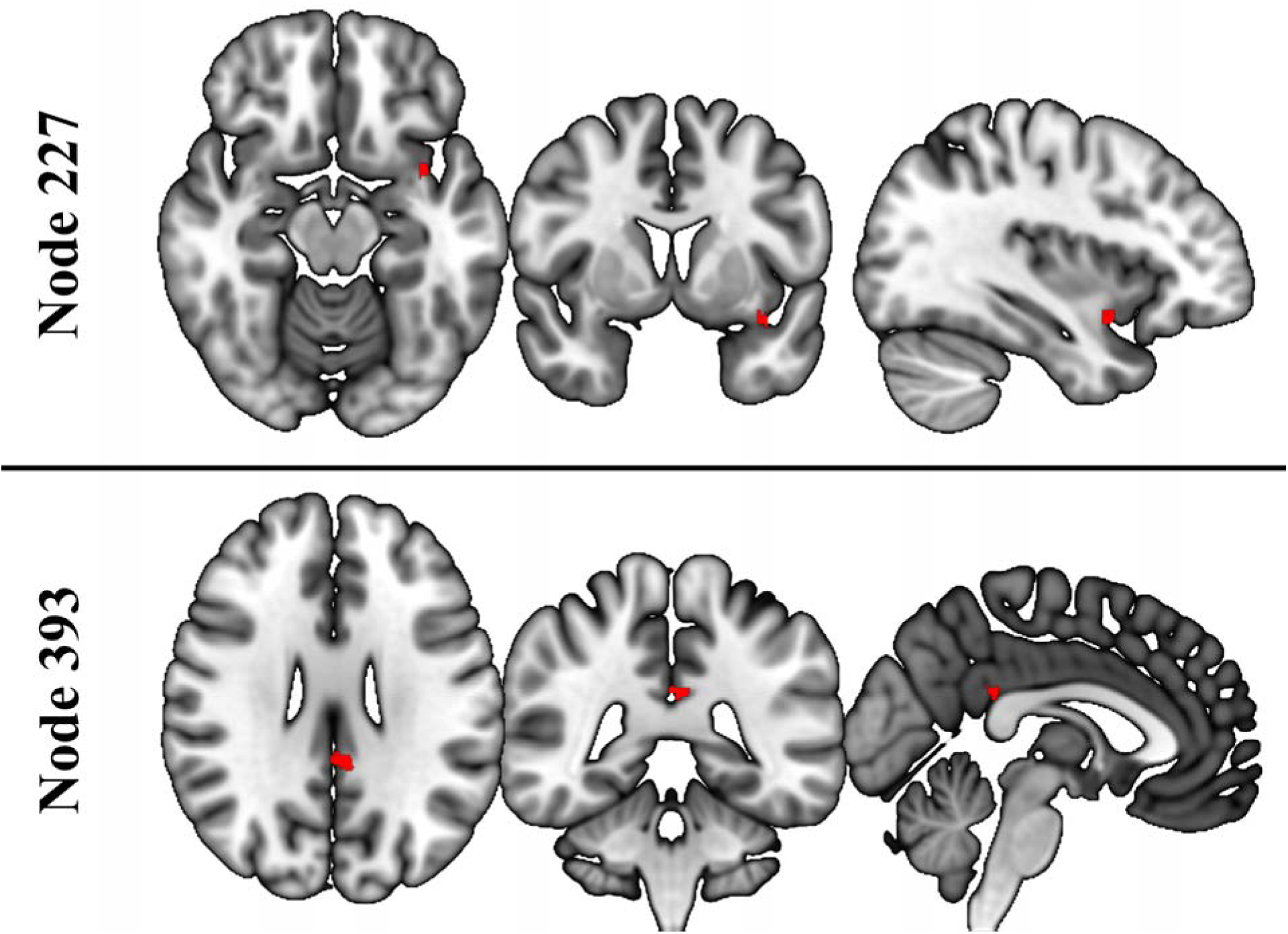
Atlas nodes lost in reslicing of the Schaefer 1000-node (17 Subnetwork) atlas. Nodes are highlighted in red. Node 227 was lost in reslicing to 4x4x4mm (for MSC analyses) and 4x4x5 mm (for NCANDA analyses). Node 393 was retained in reslicing to 4x4x4mm but lost in reslicing to 4x4x5mm. Images are in radiological orientation.

**Supplementary Figure 2.**
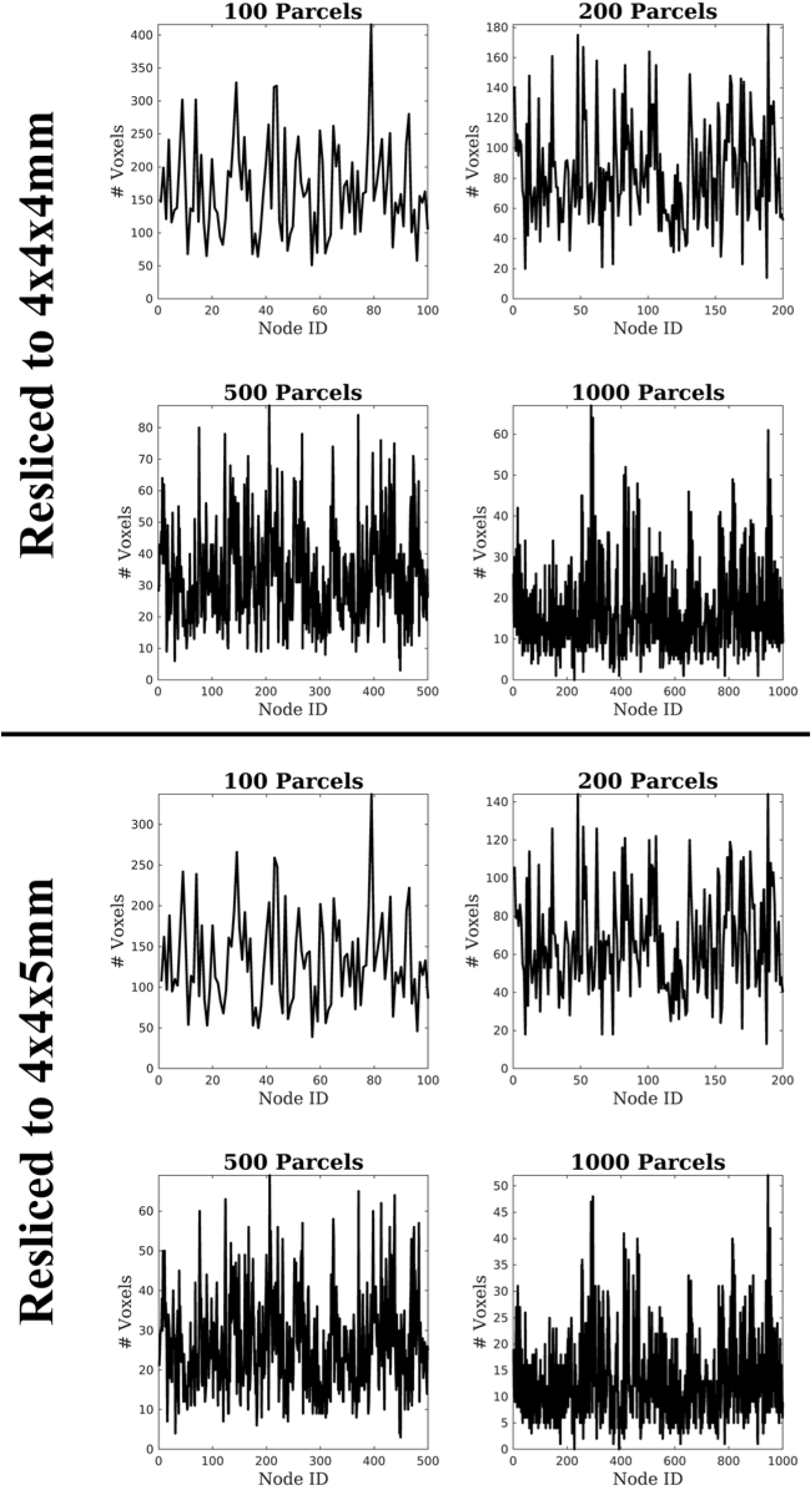
The number of voxels in each node of the Schaefer atlases is shown for both the resliced 4x4x4mm atlas and the 4x4x5mm atlas.

**Supplementary Figure 3.**
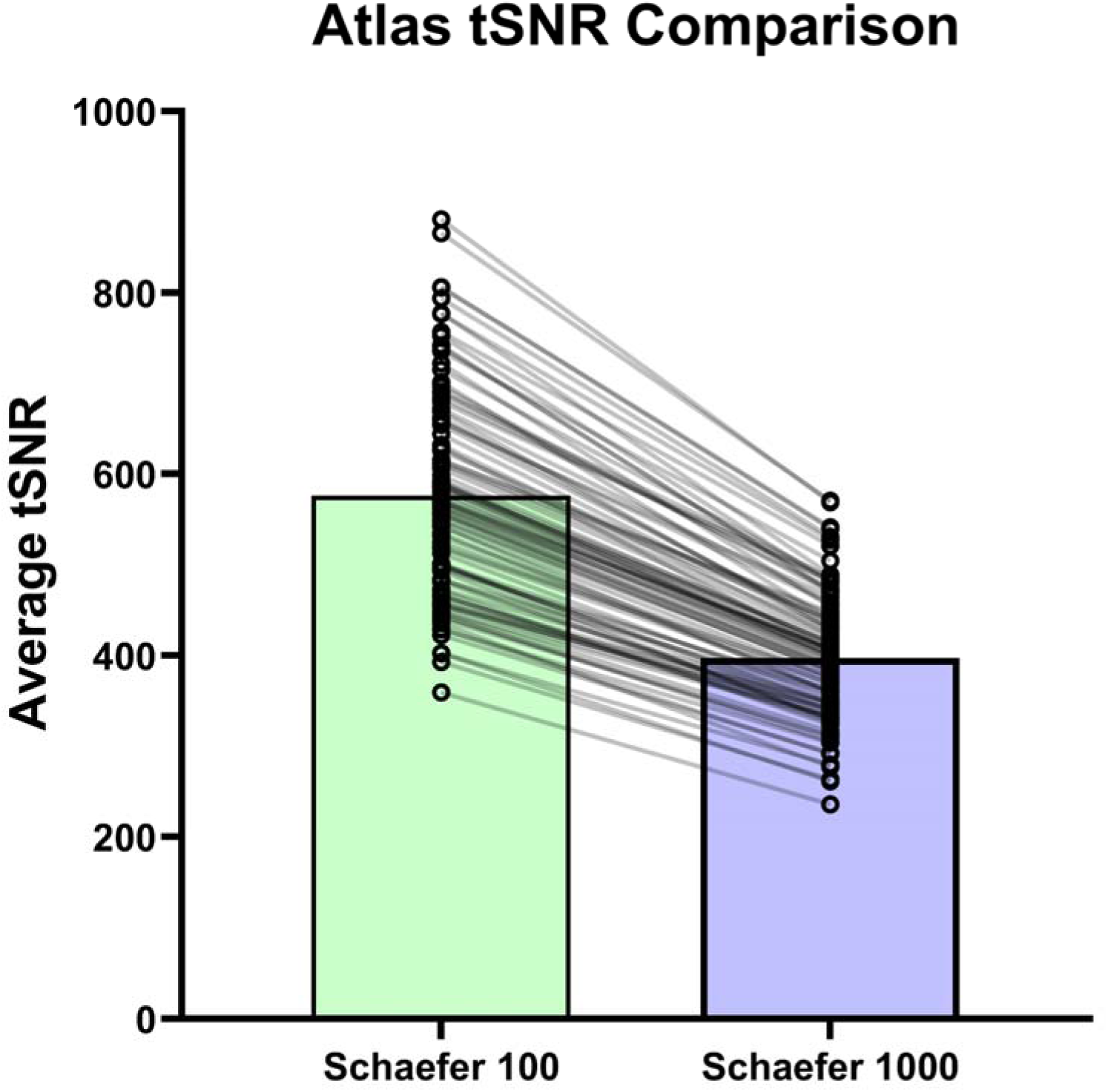
Comparison of tSNR (averaged across all nodes) for MSC scans parcellated into the Schaefer 100- versus Schaefer 1000-region atlas. Each dot represents a different person’s scan. Lines connecting the two columns indicate the same scan. Lines are semi-transparent to help with following individual lines.

**Supplementary Table 1.**
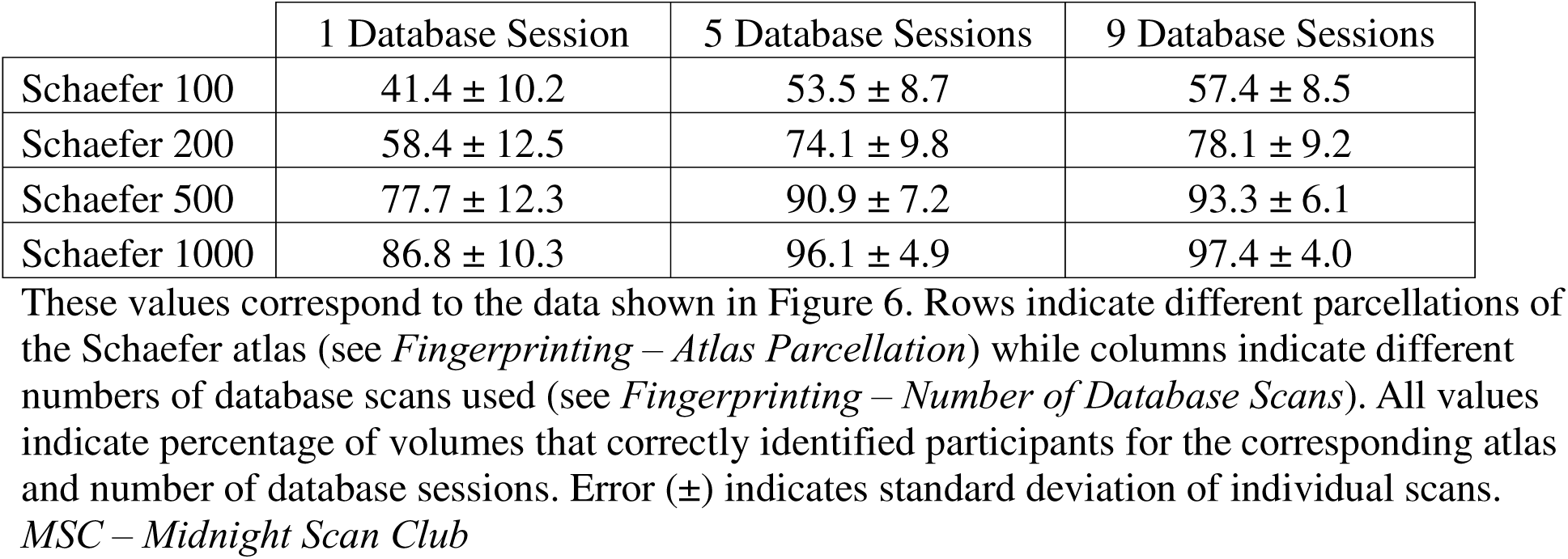
MSC Individual Volume Identification Accuracy.

**Supplementary Table 2.** MSC Full Scan Identification Accuracy.

|  | 1 Database Session | 5 Database Sessions | 9 Database Sessions |
| --- | --- | --- | --- |
| Schaefer 100 | 98.7 | 99.8 | 100 |
| Schaefer 200 | 99.9 | 100 | 100 |
| Schaefer 500 | 100 | 100 | 100 |
| Schaefer 1000 | 100 | 100 | 100 |

*MSC – Midnight Scan Club*

**Supplementary Table 3.**
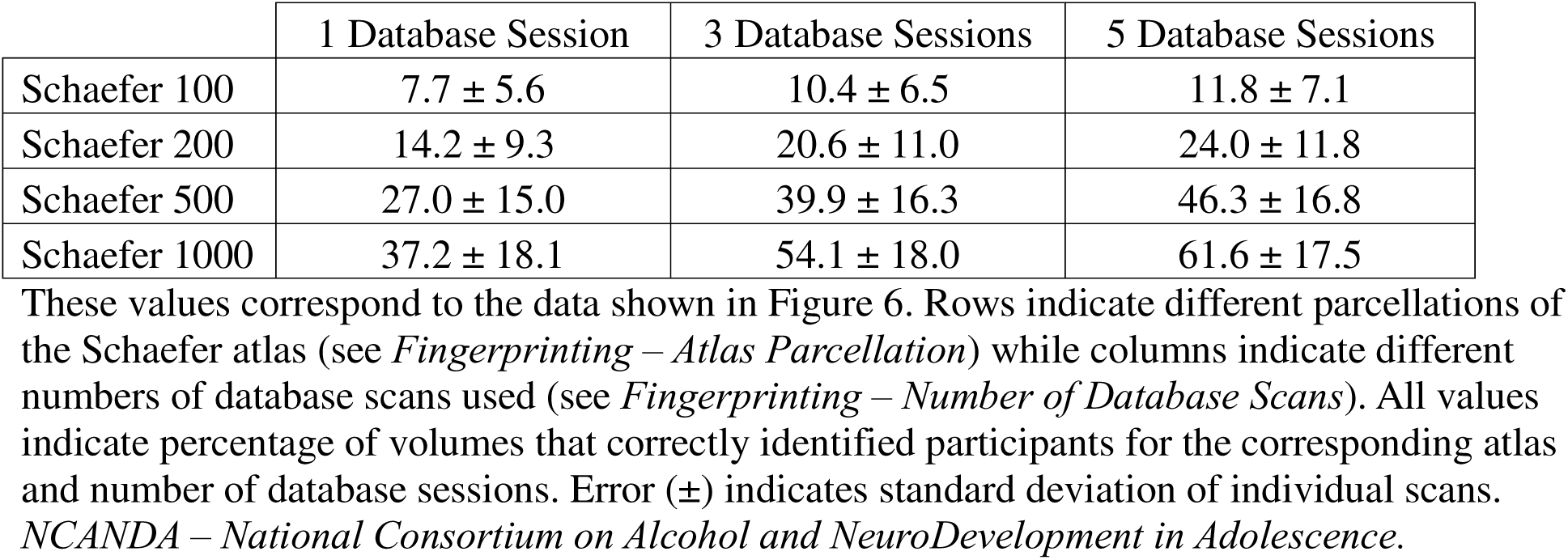
NCANDA Individual Volume Identification Accuracy.

**Supplementary Table 4.** NCANDA Full Scan Identification Accuracy.

|  | 1 Database Session | 3 Database Sessions | 5 Database Sessions |
| --- | --- | --- | --- |
| Schaefer 100 | 59.0 | 81.8 | 89.3 |
| Schaefer 200 | 80.1 | 95.8 | 97.7 |
| Schaefer 500 | 93.2 | 98.9 | 99.7 |
| Schaefer 1000 | 95.7 | 99.6 | 99.8 |

**Supplementary Table 5.** BNET Individual Volume Identification Accuracy.

|  | 1 Database Session |
| --- | --- |
| Schaefer 100 | 6.8 ± 6.4 |
| Schaefer 200 | 12.2 ± 10.5 |
| Schaefer 500 | 22.0 ± 16.8 |
| Schaefer 1000 | 30.6 ± 21.2 |

**Supplementary Table 6.** BNET Full Scan Identification Accuracy.

|  | 1 Database Session |
| --- | --- |
| Schaefer 100 | 42.8 |
| Schaefer 200 | 63.0 |
| Schaefer 500 | 77.7 |
| Schaefer 1000 | 83.5 |

**Supplementary Table 7.** MSC Individual Volume Identification Accuracy – No Gray Matter Regression.

|  | 1 Database Session | 5 Database Sessions | 9 Database Sessions |
| --- | --- | --- | --- |
| Schaefer 100 | 42.1 ± 7.7 | 54.3 ± 9.0 | 58.1 ± 9.4 |
| Schaefer 1000 | 86.8 ± 8.3 | 96.3 ± 4.9 | 97.5 ± 4.0 |

Rows indicate different parcellations of the Schaefer atlas (see *Fingerprinting – Atlas* *Parcellation*) while columns indicate different numbers of database scans used (see *Fingerprinting – Number of Database Scans*). All values indicate percentage of volumes that correctly identified participants for the corresponding atlas and number of database sessions. Error (±) indicates standard deviation of individual scans. *MSC – Midnight Scan Club*

**Supplementary Table 8.** MSC Full Scan Identification Accuracy – No Gray Matter Regression.

|  | 1 Database Session | 5 Database Sessions | 9 Database Sessions |
| --- | --- | --- | --- |
| Schaefer 100 | 97.9 | 99.4 | 99.0 |
| Schaefer 1000 | 99.9 | 100 | 100 |

Rows indicate different parcellations of the Schaefer atlas (see *Fingerprinting – Atlas Parcellation*) while columns indicate different numbers of database scans used (see *Fingerprinting – Number of Database Scans*). All values indicate percentage of full scans that correctly identified participants across all iterations of different target scans and subjects.
*MSC – Midnight Scan Club*

**Supplementary Table 9.** MSC Individual Volume Identification Accuracy – High/Low Motion.

|  | High Motion | Low Motion |
| --- | --- | --- |
| Schaefer 100 | 37.7 | 57.5 |
| Schaefer 200 | 64.1 | 78.1 |
| Schaefer 500 | 85.2 | 93.4 |
| Schaefer 1000 | 88.8 | 97.4 |

Rows indicate different parcellations of the Schaefer atlas (see *Fingerprinting – Atlas Parcellation*) while columns indicate volumes with high motion versus low motion. Accuracy is based on fingerprinting analyses with 9 database scans. All values indicate percentage of full scans that correctly identified participants across all iterations of different target scans and subjects.
*MSC – Midnight Scan Club*

**Supplementary Table 10.** NCANDA Individual Volume Identification Accuracy – High/Low Motion.

|  | High Motion | Low Motion |
| --- | --- | --- |
| Schaefer 100 | 7.3 | 12.0 |
| Schaefer 200 | 15.0 | 24.4 |
| Schaefer 500 | 30.3 | 46.9 |
| Schaefer 1000 | 42.8 | 62.3 |

Rows indicate different parcellations of the Schaefer atlas (see *Fingerprinting – Atlas Parcellation*) while columns indicate volumes with high motion versus low motion. Accuracy is based on fingerprinting analyses with 5 database scans. All values indicate percentage of full scans that correctly identified participants across all iterations of different target scans and subjects.
*NCANDA – National Consortium on Alcohol and NeuroDevelopment in Adolescence*

**Supplementary Table 11.** BNET Individual Volume Identification Accuracy – High/Low Motion.

|  | High Motion | Low Motion |
| --- | --- | --- |
| Schaefer 100 | 5.2 | 6.8 |
| Schaefer 200 | 9.0 | 12.4 |
| Schaefer 500 | 17.8 | 22.2 |
| Schaefer 1000 | 23.5 | 31.0 |

Rows indicate different parcellations of the Schaefer atlas (see *Fingerprinting – Atlas Parcellation*) while columns indicate volumes with high motion versus low motion. Accuracy is based on fingerprinting analyses with 1 database scan. All values indicate percentage of full scans that correctly identified participants across all iterations of different target scans and subjects.
*BNET – Brain Networks and Mobility*

**Supplementary Table 12.** MSC Static Network Identification Accuracy.

|  | 1 Database Session | 5 Database Sessions | 9 Database Sessions |
| --- | --- | --- | --- |
| Schaefer 100 | 96.1 | 99.6 | 100 |
| Schaefer 200 | 99.0 | 100 | 100 |
| Schaefer 500 | 100 | 100 | 100 |
| Schaefer 1000 | 99.9 | 100 | 100 |

Percentage of scans for which the individual was correctly identified based on their static network. Rows indicate different parcellations of the Schaefer atlas (see *Fingerprinting – Atlas Parcellation*) while columns indicate different numbers of database scans used (see *Fingerprinting – Number of Database Scans*). All values indicate percentage of full scans that correctly identified participants across all iterations of different target scans and subjects.
*MSC – Midnight Scan Club*

**Supplementary Table 13.** NCANDA Static Network Identification Accuracy.

|  | 1 Database Session | 3 Database Sessions | 5 Database Sessions |
| --- | --- | --- | --- |
| Schaefer 100 | 71.7 | 89.7 | 93.5 |
| Schaefer 200 | 83.4 | 95.3 | 97.7 |
| Schaefer 500 | 90.6 | 97.6 | 98.5 |
| Schaefer 1000 | 92.4 | 98.7 | 99.3 |

Percentage of scans for which the individual was correctly identified based on their static network. Rows indicate different parcellations of the Schaefer atlas (see *Fingerprinting – Atlas Parcellation*) while columns indicate different numbers of database scans used (see *Fingerprinting – Number of Database Scans*). All values indicate percentage of full scans that correctly identified participants across all iterations of different target scans and subjects.
*NCANDA – National Consortium on Alcohol and NeuroDevelopment in Adolescence*

**Supplementary Table 14.** BNET Static Network Identification Accuracy.

|  | 1 Database Session |
| --- | --- |
| Schaefer 100 | 48.3 |
| Schaefer 200 | 61.0 |
| Schaefer 500 | 72.5 |
| Schaefer 1000 | 76.3 |

Percentage of scans for which the individual was correctly identified based on their static network. Rows indicate different parcellations of the Schaefer atlas (see *Fingerprinting – Atlas Parcellation*). All values indicate percentage of full scans that correctly identified participants across all iterations of different target scans and subjects.
*BNET – Brain Networks and Mobility*

**Supplementary Table 15.** Midnight Scan Club Task Identification.

|  | Faces | Words | Rest |
| --- | --- | --- | --- |
| **Within Subject** | 58.4 ± 10.6 | 46.4 ± 10.3 | 60.3 ± 14.4 |
| **Between Subjects** | 49.9 ± 9.0 | 32.1 ± 6.4 | 51.4 ± 8.5 |

## Notes

### Competing Interest Statement

The authors have declared no competing interest.

